# MYC and Epithelial to Mesenchymal Transition (EMT) Independently Predict Circadian Rhythm Disruption in Lung Adenocarcinoma

**DOI:** 10.1101/2025.08.15.670530

**Authors:** Jamison B. Burchett, Aslihan Ambeskovic, McKayla Ford, Jacob Cody Naccarato, Juliana Cazarin, Fabio Hecht, Molly Hulver, Xueyang He, Joshua C. Munger, Paula M. Vertino, Isaac S. Harris, Stephano S. Mello, Brian J. Altman

## Abstract

The molecular circadian clock is known to be disrupted in lung adenocarcinoma, and its disruption is pro-tumorigenic in mouse models of this disease. However, the determinants of disruption of the molecular clock in human cancer are not clear. We hypothesized that derangement in expression of specific circadian clock genes or elevated MYC expression could correlate with circadian disruption in human tumors, and used Clock Correlation Distance (CCD) to compare clock order and strength in tumors based on the expression of these genes. While the expression of individual circadian genes did not consistently correlate with disruption, tumors with the highest expression of MYC or high MYC pathway activation had significantly disrupted rhythms compared to those with lower MYC. Unexpectedly, a subset of tumors with very low levels of MYC, below that found in normal lung, also showed disruption of circadian rhythms, prompting us to explore novel determinants of disruption in these tumors. We found that expression of programs associated with epithelial to mesenchymal Transition (EMT) and TGF-β signaling were enriched in tumors with the lowest MYC expression, and that, surprisingly, those tumors with a mesenchymal expression pattern had more ordered (stronger) rhythms. To directly test this correlation between cell state and rhythms, we exposed lung adenocarcinoma cells to TGF-β to induce EMT. TGF-β induced a quasi-mesenchymal phenotype and caused a significant increase in the amplitude of oscillations in these cells. Together, our data show that MYC expression, pathway activation, and a mesenchymal cell state are both independent determinants of circadian status in lung adenocarcinoma.

## Introduction

The twenty-four-hour cycle known as circadian rhythm helps to regulate several core biological processes in cells such as cell division and metabolism (1–6). Rhythms are synchronized by external stimuli such as light exposure, which is processed by the suprachiasmatic nucleus within the brain, and signals are sent out to the rest of the body to entrain cells in different organs to the same internal phases. Each cell has its own individual molecular clock that responds to these signals, with core clock transcription factors BMAL1 and CLOCK regulating expression of their target genes and making up the positive arm of the clock. Some of their targets include the REV-ERB, CRY, and PER protein families that inhibit the expression and function of the CLOCK and BMAL1 heterodimer, leading to negative feedback, and are the negative arms of the clock. This system of feedback loops leads to the rhythmic expression of hundreds to thousands of ‘output’ genes in each cell type (7,8), and creates the twenty-four-hour cycle that we observe as circadian rhythms within cells.

Disruption of the molecular clock is widespread in human cancer (9,10). However, one of the difficulties in studying the interactions between circadian rhythms and cancer is that there is not currently a way to determine which factors may be associated with clock disruption in human tumors. Several methods were developed to use single time point expression data from patient tumors, one of which is Clock Correlation Distance (CCD) (9). CCD is a computational method that compares the relationships between a set of circadian genes to a reference correlation made from multiple healthy mouse tissue types. It was validated on normal healthy human tissue and was used to demonstrate disorder in circadian rhythms in various cancer types including breast, colon, and lung adenocarcinoma. Other computational methods have been published that agree with the findings of CCD (10–12). Supporting the potential clinical importance of circadian rhythm disruption in human tumors, genetic and behavioral disruption of circadian rhythms was shown to accelerate tumorigenesis in mouse models of neuroblastoma, melanoma, liver, lung, pancreatic, and colorectal cancer (3,4,13–16).

Circadian disorder as a potential driver of tumorigenesis is not universal across all cancer types, as some cancer types still have functioning clocks within their tumors(17–21). For this study, we focused on non-small cell lung cancer, and specifically lung adenocarcinoma (LUAD). The lung is a highly circadian organ, and in mice, the lung has the third highest number of oscillating transcripts in the body (7). The American Cancer Society reported that in 2024 there were 234,580 new cases of lung cancer with 125,070 deaths making it the third most common cancer and the leading cause of cancer deaths in the United States. Circadian disruption is known to accelerate lung adenocarcinoma formation and decrease survival in mouse models but mutations and deletions in circadian genes are not normally observed in human lung cancer. Instead, there are changes to the regulation and expression of both core and peripheral circadian genes observed in lung cancer as well as other cancer types (9,11,12,14,16). This makes lung adenocarcinoma a prime target for the further study of factors that may lead to circadian disruption in cancer.

One of the mechanisms that the circadian clock is disrupted in cancer is through overexpression of the MYC oncoprotein (13). The MYC family, which is amplified or translocated in ∼30% of human cancers and widely amplified in human lung cancer (22), can disrupt the circadian clock through multiple pathways including upregulation of negative-arm factors such as REV-ERB α and β and direct suppression of CLOCK and BMAL1, causing the clock to stop oscillating (13,23). However, it is likely that elevated MYC only accounts for a portion of the widespread clock disruption observed in multiple human cancer types, and other factors and pathways likely contribute as well. Additionally, the role of MYC in potential circadian rhythm disruption in lung adenocarcinoma has not yet been investigated, nor have other potential determinants of circadian disruption in this disease been elucidated.

In this study, we used tumor expression data from lung adenocarcinoma samples from The Cancer Genome Atlas (TCGA) to investigate what genes and pathways correlate with circadian disruption. We began by assessing whether variations in individual molecular clock genes could be viable markers of circadian disruption and found that expression of individual circadian genes did not consistently correlate with circadian disruption in LUAD but considering them as a pathway showed some potential correlation with circadian status. In contrast, both high expression of MYC and high MYC pathway activation correlated with circadian disruption. Interestingly, tumors with very low MYC (below the level observed in normal lung) also showed disruption of rhythms, indicating a potential MYC-independent factor controlling the strength of the clock in these tumors. We compared these tumors with those that had the strongest rhythms and discovered that epithelial to mesenchymal transition (EMT) and TGF-β signaling pathways were independent markers of circadian disruption, and that the tumors with very low MYC and disrupted rhythms were more epithelial in nature and had less evidence of TGF-β signaling. To test if this computational finding could be observed *in vitro*, we exposed mouse lung cancer cell lines that have been characterized as more epithelial to TGF-β to induce EMT and observed a significant increase in the amplitude (strength) of their circadian oscillations compared to control cells. These results indicate that enrichment in the MYC, EMT, and TGF-β pathways could all be used to infer and predict circadian status in lung adenocarcinoma tumors.

## Methods

### Data Acquisition

Gene expression data was downloaded from the National Cancer Institutes GDC Data portal. In the Project tab of the portal, projects were filtered for Bronchus and Lung primary site that were adenomas and adenocarcinomas, and the selected projects were CPTAC-3 (dbGaP accession number phs001287), TCGA-LUAD (dbGaP accession <u>phs000178.v11.p8</u>) , and TCGA-LUSC (dbGaP accession <u>phs000178.v11.p8</u>) which were then saved as a cohort within the portal. This cohort was then used in the Repository program of the portal, and open access RNA-seq data from primary solid tissue tumor samples as well as associated normal samples were selected for download. Sample files were then downloaded along with associated Clinical and Metadata files, and all sample files were compiled into a single expression dataset by creating a list of all sample expression file names and then performing a left join using the list of sample file names and expression files, creating a single file with labelled samples and expression data. This process was also repeated to create a metadata file with containing all labelled samples and reported clinical data.

### Determining sample groups

Data samples were sorted into quartiles based on individual expression or enrichment scores of the specified target for each individual analysis. The data was sorted in R version 3.4.3 using the dplyr function ntile v1.1.4 which sorts samples into equal sized bins based on the specified value for each sample. In cases where the total number of samples is not divisible by 4, the extra samples were placed into the earlier bins.

### Visualization of survival by gene expression

Kaplan-Meier (KM) plots were created using KMPlot.com (24,25) using aggregated data from several mRNA gene chip lung cancer databases. Individual genes correlation to survival were compared by the highest and lowest tertiles of expression in lung adenocarcinoma samples, with a p-value of <0.05 indicating a significant difference in survival between the two groups.

### Determination of circadian disorder by CCD

The Clock Correlation Distance analysis, R package deltaCCD, v1.0.4, described previously (9), was used to compare sorted sample groups of human lung samples to the established reference clock to determine CCD scores for each analyzed group. Because deltaCCD (which compares CCD scores to each other to assess significance) is a one-sided analysis, in each analysis the group with the lowest reported CCD score, representing the group with a clock most similar to the reference, was identified as the control group for further comparisons. deltaCCD was then calculated comparing all other groups to the control group with a p-value of <0.05 indicating a significant difference between groups. The deltaCCD calculation determines significance through 1000 rounds of label permutation, keeping the reference samples constant.

### Differential Gene Expression Analysis by cBioportal

Data from the Lung Adenocarcinoma (TCGA, Pan Cancer Atlas) Study in cBioportal (26–28) was used for these analyses. Custom groups were created and labeled using relevant sample IDs for each group determined by previous quartile analysis described above for groups of interest (i.e. MYC and MYC_SUM expression quartiles). Once groups were created comparisons were performed within the cBioportal groups tab by selecting the desired sample groups. Upon comparing the desired groups, we navigated to the mRNA tab of the comparison which reports differentially expressed genes and filtered the list to only significantly differentially expressed genes, q-value <0.05 derived from a Benjamini-Hochberg procedure within cBioportal. We then sorted them by their Log2 ratio before downloading the gene list.

### Differential Pathway Enrichment analysis by GSEA

Gene Set Enrichment Analysis has been previously described (29,30). We used GSEA on a pre-ranked gene list produced by the group comparison in cBioportal previously detailed. For each database compared using the pre-ranked gene list, we chose no collapse/remap to gene symbols for the analysis with 1000 permutations. Database from msigdb (31–33) used were: C2.biocarta, C2.pid, C2.reactome, C2.wikipathways, C2.all, C2.cpg, C2.cp, C2.KEGG, C3.all, C3.tft, C3.tft.gtrd, C3.tft.legacy, C5.all, C5.go, C5.go.bp, C5.go.cc, C5.go.hpo, C5.go.mf, C6.all, C7.all, C7.immunesigdb, C7.vax, C8.all, and h.all (Hallmarks).

### Determination of individual sample pathway enrichment using GSVA

Gene Set Variation Analysis has been previously described (34) and the GSVA R package version 1.34.0 was used with default parameters to apply individual relative enrichment scores for each sample for each given gene set. Gene lists were identified by either the genes that make up GSEA pathways, or previously established gene list of circadian genes (35). All code used for GSVA analysis is available on FigShare (see below).

### Assignment of cellularity scores

Determination of cellularity scores, a measure of tumor purity, is described previously (36). This study established cellularity scores for samples across several cancer types in the cancer genome atlas through multiple methods including a median purity level from normalizing all other methods.

This median measure reported as the consensus measurement of purity (CPE) was used as our representation of tumor purity. Values were downloaded and matched to our sample ID’s, appending the CPE values onto the existing metadata for each sample that had a calculated score. Samples that did not have a reported CPE value were not included in the following cellularity group analyses.

### Cell Culture

CMT64 were purchased from Sigma Aldritch from their ECACC (European Collection of Authenticated Cell Cultures) collection (Sigma, St. Louis, MO, USA). Cells were cultured in Dulbecco’s Modification of Eagle’s Medium (DMEM) (Corning, Manassas, VA, USA, Catalog # 10-013-CV) with 10% Fetal Bovine Serum (FBS) (GeminiBio, West Sacramento, CA, USA, Catalog# 15140-122) and 1% of Penicillin Streptomycin (PS) (Gibco, Grand Island, NY, USA, Catalog# 100-106). A549 and H1299 cell lines were cultured in RPMI 1640 with 10% FBS and 1% PS. A549 and H1299 were a generous gift of Dr. Paula Vertino (University of Rochester, Rochester, NY, USA) and were authenticated with the IDEXX Cell Check 16 service (IDEXX, Columbia, MO, USA). All cell lines were maintained in humidified tissue culture incubators kept at 5% CO_2_ and 37°C.

### Establishing MYC-ER Cells

The C-MYC-ER^TM^ plasmid, described previously (37), used as an inducible form of MYC overexpression responds to 4-hydroxytamoxifen and is in a p-Babe-Zeocin resistant vector (13). Viral supernatant was produced by 293T cells using the pCMV-VSVG and pUMVC vectors. Viral supernatant was filtered through a 0.45um filter prior to centrifugation in a 10,000MW centrifuge filter (Cytiva, Wilmington, DE, USA, Catalog # 28932360). The resulting viral liquid was aliquoted and either used immediately for infection of cells or snap frozen in liquid nitrogen and kept at −80°C. Infected H1299 and A549 cells were selected with 50µg/mL Zeocin (Invitrogen, Carlsbad, CA, USA, Catalog# R25001) and treated with 0.5 µM 4-hydroxytamoxifen to induce translocation of the MYC-ER construct to the nucleus.

### Protein Harvest and Western Blotting

Protein was extracted by lysing cells using the M-Per lysis (Thermo Fisher Scientific, Rockford, IL, USA, Catalog #78501) reagent combined with protease and phosphatase inhibitor cocktail (Sigma-Aldritch, St. Louis, MO, USA, Catalog #PPC1010) and phosphatase inhibitor cocktail 2 (Sigma-Aldritch, St. Louis, MO, USA, Catalog # P52726). Cells were incubated in the lysis mixture on ice for twenty minutes followed by a spin at max speed (13,000 xg) in a centrifuge pre-cooled to 4°C for ten minutes. Following this spin, supernatant was collected to be used for western blot analysis or stored at −80°C for later use. Protein samples were quantified using the Bio-Rad DC Protein Assay. For the western blot, aliquots of equal concentration of protein were made and a 5x SDS loading dye/5% β-mercaptoethanol solution was added. The samples were then incubated at 99°C for 10 minutes followed by a 1 minute spin at 13,000 rcf before being loaded into a Bio-Rad Criterion 4-20% TGX Stain-Free precast gel (Bio-Rad, Hercules, CA, USA) and ran using a Bio-Rad PowerPac Basic at 140V with 1x Tris/Glycine/SDS Buffer (Bio-Rad, Hercules, CA, USA, Catalog #1610732) until the dye had run off the gel, approximately 1 hour. The gels were then transferred to a nitrocellulose membrane(Bio-Rad, Hercules, CA, USA, Catalog #1620167) using the Trans-Blot Turbo system (Bio-Rad, Hercules, CA, USA). Stain free blots were then imaged using the ChemiDoc MP imager (Bio-Rad, Hercules, CA, USA) using the auto-exposure setting for a total protein image used for normalization. In **Figure 3**, total protein is collapsed to the size of a single band for ease of viewing. The following primary antibodies were used: rabbit anti-E-cadherin (Cell Signaling, MA, USA, Catalog #3195), rabbit anti-C-MYC (Abcam, Waltham, MA, USA Catalog #AB32072), and rabbit anti-BMAL1 (Cell Signaling, MA, USA, Catalog #14020). The following secondary antibody was used: goat anti-rabbit Alexa Fluor 680 (Thermo Fisher Scientific, Rockford, IL, USA, Catalog #A11357). Stained blots were imaged using the Chemi-Doc MP imager using the auto-exposure setting and images were analyzed and quantified using the Bio-Rad Image Lab software v6.0.1. Band intensity was calculated minus background and normalized using total protein for comparisons.

### Imaging (Spark Cyto)

CMT64 cells were plated in 12-well tissue culture plates at 10,000 cells per well in 1mL of media. Following plating, cells were incubated for 24 hours before being exposed to 5ng/mL of TGF-β. The plate was then imaged using the Tecan Spark Cyto imager at 24, 48, and 72 hours post exposure. The plate was imaged with media and kept at 37°C and 5% CO_2_ for the duration of the imaging. The Spark Cyto program performed brightfield 2D imaging with the 4x objective for all wells and runs of the plate and performed counting and area analysis on all images reporting total cell counts, percent confluence, and roughness measurements for each well.

### RNA Extraction and Quantitative PCR

All RNA was extracted using the E.Z.N.A HP Total RNA (Omega Bio-tek, Norcross, GA, USA, Catalog # R6812-02) following the kits provided protocol. RNA was then used to make complimentary DNA using TaqMan Reverse Transcription Reagents (Applied Biosystems, Carlsbad, CA, USA, Catalog #N8080234), and qPCR was performed using PerfeCTa SYBR Green Fastmix (QuantBio, Beverly, MA, USA, Catalog #95074-05K) with Quant Studio 5 quantitative PCR machines from Applied Biosystems. Samples were set up with triplicate technical repeats, and relative expression was normalized to β2M and calculated by the ΔΔCT.

### Monitoring changes in CMT64 BMAL1 oscillations by Lumicycle analysis

CMT64 cells with a BMAL1::LUC construct (38) expressed were plated in 35mm tissue culture dishes at 400,000 cells per dish in 2mL of complete DMEM media in triplicate per condition. Twenty-four hours after plating the plates were exposed to either 5ng/mL TGF-β or vehicle. Seventy-two hours post treatment, all plates were exposed to 0.1µM dexamethasone to synchronize the cells for two hours. Following the synchronization period, the media was aspirated and replaced with phenol-free DMEM, 10% FBS, 1% PS, 1M HEPES, and 4mM L-glutamine with 0.1mM beetle luciferase (Promega, Madison, WI, USA, Catalog # E1602). The plates were then sealed with vacuum grease (High Vacuum Grease, Molykote, Wilmington, DE, USA, Catalog #2021854-0221) surrounding the lids to prevent evaporation while in the Lumicycle instrument (Actimetrics, Wilmette, IL, USA) and placed into the Lumicycle at atmospheric CO_2_ and 37°C to have their luminescence measured every 30 minutes for 4 days. Results were analyzed using the Lumicycle Analysis software v2.701 (Actimetrics Software), and raw data was input into the analysis software which then determined a baseline and calculated a baseline-subtracted data series. This baseline subtracted data was then fit to a damped sin wave to calculate average amplitude for each sample. Oscillations and calculated wave amplitude were averaged across triplicates with standard errors of the mean calculated and a two-sided Welch’s T-Test performed to determine significance.

### Data and code availability

All raw data, processed data, and code used in this manuscript is available on FigShare at https://doi.org/10.60593/ur.d.c.7909358. This link will be made ‘live’ upon acceptance.

## Results

### Non-Small Cell Lung Cancer samples have significant circadian disorder, and circadian gene expression correlates with survival

We first asked whether the molecular circadian clocks of NSCLC samples were significantly more disordered than normal lung tissue samples, as was previously shown in individual subtypes of NSCLC(9). To accomplish this, we compiled gene expression data from The Cancer Genome Atlas (TCGA) and Clinical Proteomic Tumor Analysis Consortium (CPTAC) for lung adenocarcinoma and squamous cell carcinoma, N = 1240, and associated normal samples, N = 296, for comparison. We then used Clock Correlation Distance (CCD) to compare the bulk NSCLC samples to normal (**Figure 1A**). CCD is a computational method that compares the relationships between a set of circadian genes to a reference correlation made from multiple healthy mouse tissue types. The more different the relationship between the set of circadian genes is in the sample data from the reference, the higher the resulting CCD score, and the more disordered / disrupted the rhythm is. A ΔCCD can then be calculated by comparing two groups’ results to determine if the two groups’ levels of disorder are statistically significantly different. Using this method, we found that the NSCLC samples had a CCD score of 4.899 and the normal tissue samples had a score of 3.635 (**Figure 1A Right Panel**) with a ΔCCD p-value of 0.00099 indicating that the NSCLC samples had significantly more disordered rhythms than normal. For all subsequent analyses in the manuscript, 3.7 was set as the Y-axis minimum for CCD measurements since this represents “normal” circadian function in human lung tissue. With this established, we then moved towards identifying if changes in the expression of circadian genes correlated with survival in LUAD, a type of NSCLC, and how those changes may occur.

**Figure 1:**
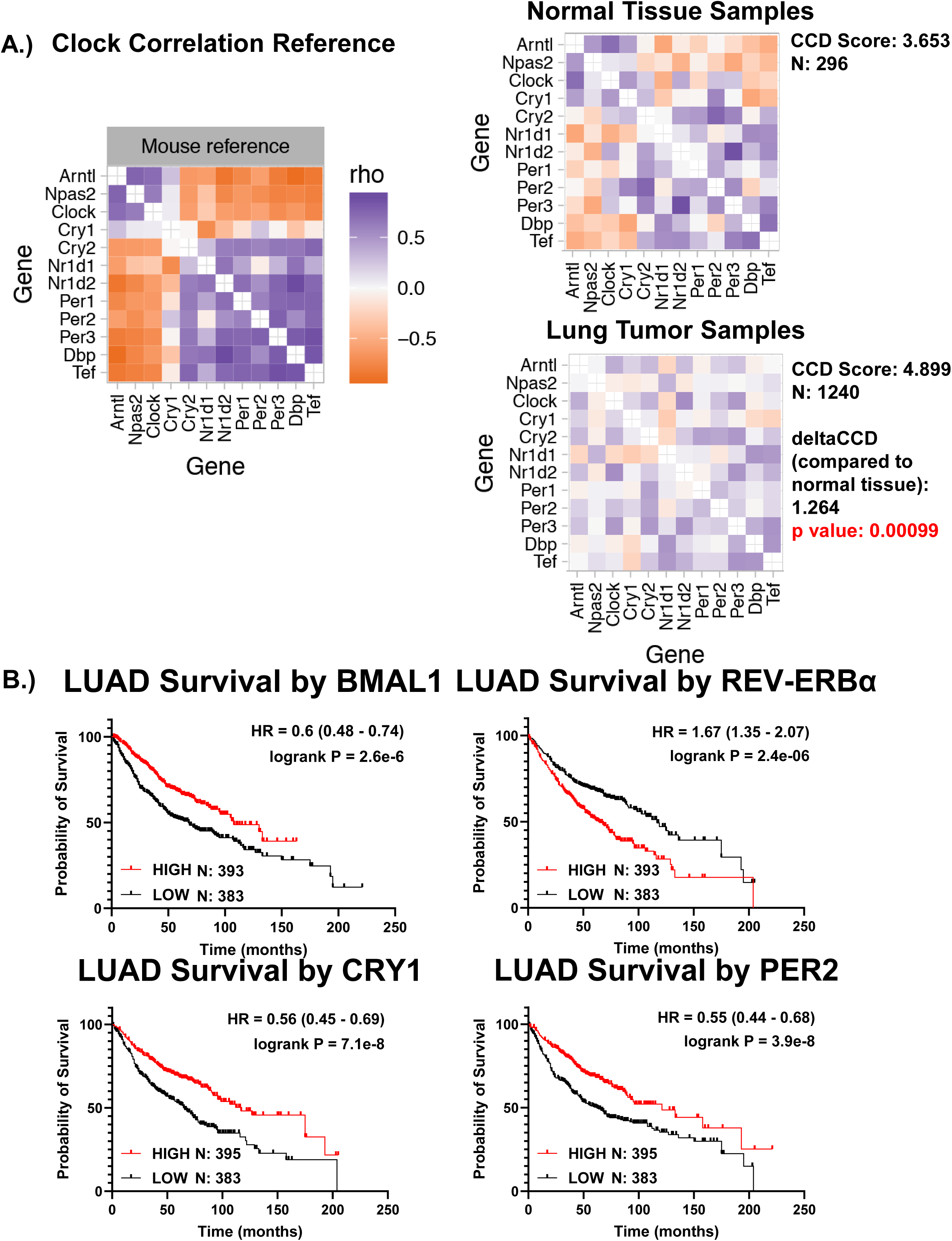
The circadian clock is disrupted in lung cancer, and molecular clock genes correlate with survival. A.) The Clock Correlation Distance (CCD) reference plot (Left) showing the spearman correlation between the core clock genes. (Right) Spearman correlation maps generated through CCD of normal lung tissue samples and lung tumor samples. Samples N’s, calculated CCD scores, deltaCCD scores, and p-values are detailed next to the correlation maps. B.) Kaplan-Meyer plots of lung adenocarcinoma patient survival comparing groups within the upper vs lower third of *BMAL1*, REV-ERBα (*NR1D1*), *PER2*, and *CRY1* expression. Plots have reported Hazard Ratio (HR), and logrank P value with a P-value <0.05 determined to be significant.

To determine whether changes in gene expression correlated with overall survival in LUAD patients, we made Kaplan-Meier plots comparing LUAD samples from a meta-analysis of several previous studies (24,25). We compared the top and bottom third of expression of individual circadian genes (*BMAL1*, *NR1D1* which encodes REV-ERBα*, PER2,* and *CRY1*) . This analysis revealed that low expression of the core circadian genes *BMAL1, Per2,* and *CRY1* and high expression of REV-ERBα were correlated with poorer survival (**Figure 1B**). Some of these patterns match the expression changes observed in previous studies of disrupted clocks in cancer published previously(13). These findings suggested that variations in circadian gene expression may impact disease progression, but the connection between these gene expression changes to the order, or disorder, of rhythms in tumors remains unclear.

### Expression of individual molecular clock genes vary significantly across LUAD, but do not consistently predict circadian disruption

We next sought to determine how variation in expression of individual molecular clock genes such as *BMAL1,* REV-ERBα (*NR1D1*), *PER2*, and *CRY1* may correlate with circadian disruption in LUAD. Since CCD compares groups of samples rather than individual samples, we began by breaking the TCGA LUAD samples into quartiles based on their expression of individual circadian genes, with the lowest expressing samples in Quartile 1 and the highest expressing samples in Quartile 4 (**Figure 2A**). For example, Quartile 1 would have the 25% of samples with the lowest *BMAL1* expression and Quartile 4 would have the 25% of samples with the highest *BMAL1* expression. We first sought to test if there was a difference in expression of these genes across the quartiles, to ensure that we were making biologically meaningful comparisons. Using this quartile method, we found that expression of *BMAL1*, REV-ERBα, *PER2*, and *CRY1* varied significantly across the quartiles (**Figure 2B**). We next performed CCD analysis on each quartile. We predicted that, due to their respective positions in the circadian clock, low BMAL1 (positive arm of the clock) and high REV-ERBα, PER2, and CRY1 (all in negative arms of the clock) would each be the most disrupted quartiles. Instead, we found that there was no significant difference in CCD score based on BMAL1 quartile and that the lowest, not highest, expressing REV-ERBα quartile was the most disrupted. While high CRY1 and PER2 quartiles were the most disrupted, the CCD scores of their quartiles did not follow a linear pattern (**Figure 2C**). We expanded our analysis to test all individual circadian genes that make up the CCD reference (**Supp.** Fig. 1A), and this unpredictability continued for each individual gene tested where its position and role in the circadian clock did not allow us to predict its impact on rhythms. Several genes tested, such as *RORC* and *PER3*, had no significant impact on disorder at all like *BMAL1*. Based on these findings, we concluded that individual circadian gene expression may not be a consistent marker to predict circadian disorder in lung adenocarcinoma.

**Figure 2:**
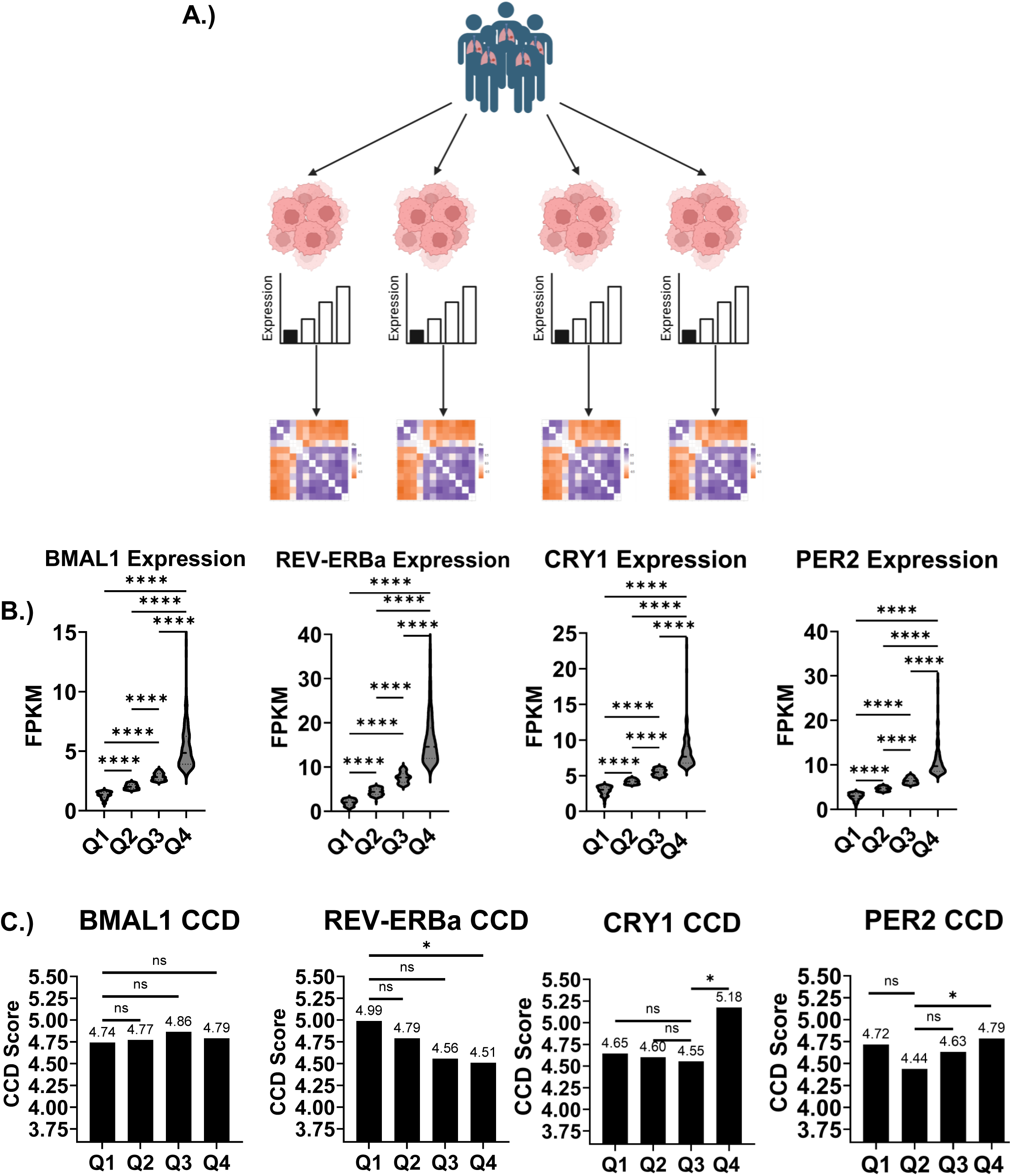
Expression of circadian genes vary across tumors, but this variation does not correlate to circadian disruption. A.) A schematic showing the breakdown of tumor samples into groups based on their expression of a target gene and then running CCD on each separate group for comparison. B.) Violin plots of expression using Fragments per kilobase per million (FPKM) values for *BMAL1*, REV-ERBα (*NR1D1*), *CRY1*, and *PER2* with median expression represented with a dotted line. Statistical difference between quartiles was determined by ordinary one-way ANOVA. **** = p<0.0001. C.) A bar chart representing the CCD scores for each quartile of expression for *BMAL1*, REV-ERBα, *PER2*, and *CRY1*. Significance was determined by sample label permutation performed by the deltaCCD package. Ns= Non-significance, * = p<0.05.

**Figure 3:**
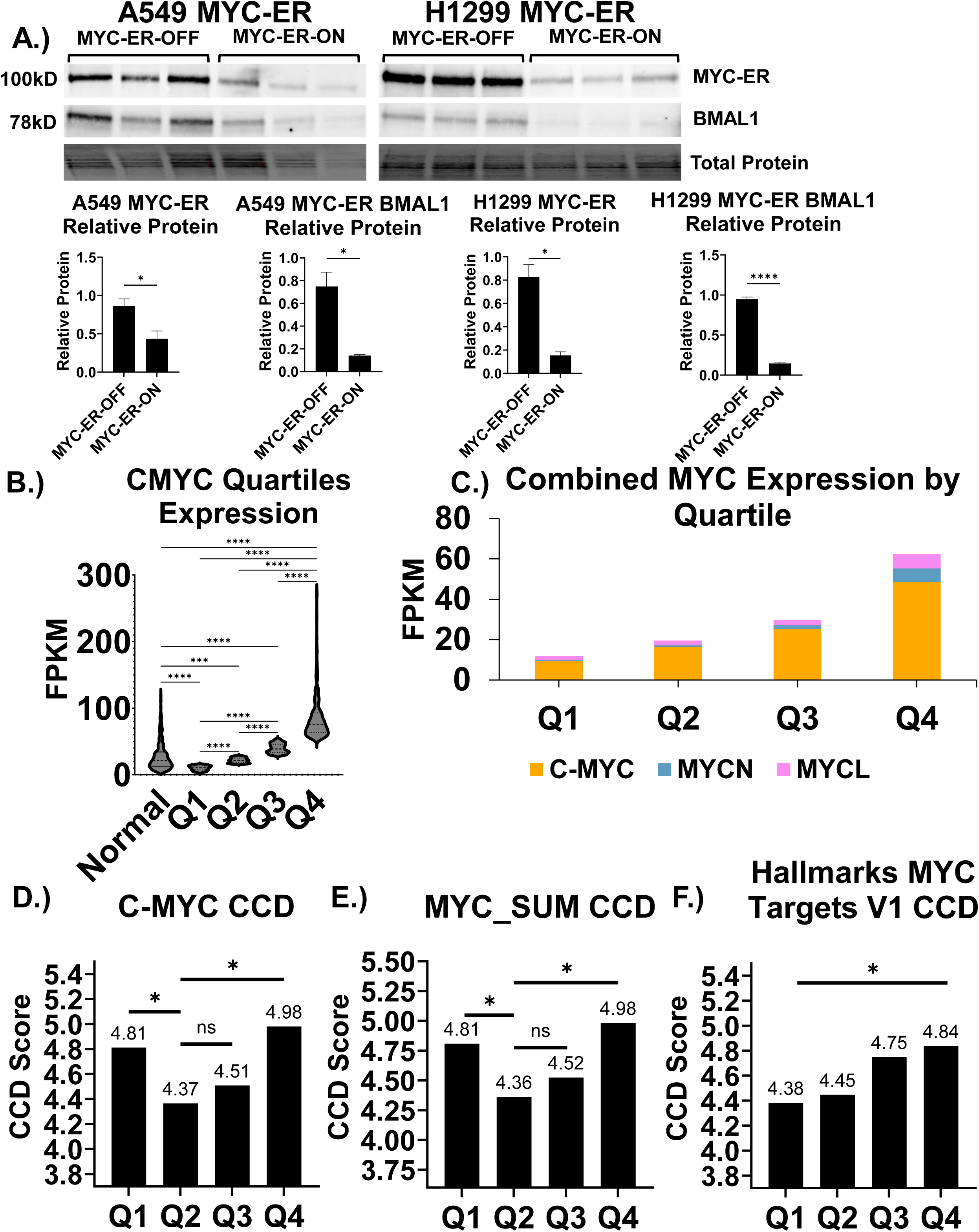
MYC oncogene expression and pathway can be used as a marker for circadian disruption. A.) Western blot of protein levels of MYC-ER and BMAL1 in human lung cancer cell lines A549 and H1299 with total protein. MYC-ER cells were treated with 4OHT (MYC-ER-ON) or EtOH (MYC-ER-OFF) for 48 hours and dexamethasone for 24 hours before protein harvest. Total protein is used as a loading control. Relative protein expression for both MYC-ER and BMAL1 in both cell lines are represented by bar charts, normalized by total protein. N = 3 biological replicates, statistical difference was determined by Welch’s T-Test. * = p<0.05, **** = p<0.0001, and data shown are representative of 3 independent experiments. B.) A violin plot of C-MYC expression using FPKM values for normal tissue samples and tumor samples previously divided into expression quartiles. Statistical difference between quartiles was determined by ordinary one-way ANOVA. **** = p<0.0001. C.) A bar chart showing the combined expression of all three types of MYC (C-, N-, and L-) in quartiles illustrating the contribution of each type of MYC to the total. D.) A bar chart representing the CCD-values of C-MYC quartiles. Significance was determined by sample label permutation performed by the deltaCCD package. Ns= non-significance, * = p<0.05. E.) A bar chart representing the CCD-values of MYC_SUM quartiles. Significance was determined by sample label permutation performed by the deltaCCD package. Ns= Non-significance, * = p<0.05. F.) A bar chart representing the CCD-values for the Hallmarks MYC Targets V1 enrichment quartiles. Significance was determined by sample label permutation performed by the deltaCCD package. * = p<0.05.

We tested if enrichment in the molecular circadian clock as a whole system could be used to predict disruption in individual samples. To acquire individual enrichment for each sample, we used Gene Set Variation Analysis (GSVA), which applies a relative enrichment score for each sample in a data set for any list of genes provided(34). We used a list of core circadian genes previously utilized by our group and others to assess clock function (35,39). GSVA allowed us to assign an individual score, representing relative enrichment of these group of genes, to each sample. We then sorted the samples into quartiles based on their enrichment in the circadian gene set and ran CCD on each of the quartiles. The P value for CCD (compared to the reference control) was not significant for quartiles 1-3, indicating a significant amount of noise or variation in these genes among the individual LUAD samples; however, there was a trend towards the lower enriched quartiles (indicating lower circadian gene expression) having a higher CCD score (more disorder) than the higher enriched quartiles (**Supp.** Fig. 1B). Overall, these results show that while individual circadian gene expression is not a consistent biomarker for circadian disorder in lung cancer, enrichment amongst a group of these genes could, with some refinement, be used as a predictor in LUAD.

### Expression of *MYC*-family genes correlates with circadian disruption in lung adenocarcinoma

We and others previously showed that MYC can alter circadian gene and protein expression in various cancer cell types (13,22,23,40,41), but it was not investigated in LUAD. To test whether MYC can induce changes in circadian protein levels in human cell line models of LUAD, we overexpressed inducible MYC-ER^TM^ in two lung adenocarcinoma cell lines that lack MYC amplification, H1299 and A549. MYC-ER^TM^ is normally inactive in the cytoplasm (MYC-ER-OFF) but translocates to the nucleus and becomes active when cells are exposed to 4-hydroxytamoxifen (MYC-ER-ON) (37). We found that when both cell lines were exposed to 4-hydroxytamoxifen MYC-ER became active, indicated by a decrease in the MYC-ER band caused by active turnover of the protein, and BMAL1 protein levels significantly decreased (**Figure 3A**). Our results, in addition to previous studies, indicating that the oncogene MYC can alter levels of multiple circadian genes and proteins, led us to test if MYC expression level could be independently used as a marker for circadian disorder. Using the same quartile strategy as we did with the individual circadian genes, we sorted the samples based on their expression of C-MYC and examined MYC expression in normal lung (**Figure 3B**). We determined that Quartiles 3 and 4 had elevated MYC compared to normal, and MYC expression significantly varied across all groups (**Figure 3B**). As C-MYC is the predominant version of MYC present in LUAD, we tested it individually as well as combining the expression of all three types of MYC: C-, N-, and L-MYC (MYC_SUM), to capture any MYC effect not entirely explained by C-MYC (**Figure 3C**). Due to the reported interactions between MYC and the circadian clock (13,22,23,40,41), we predicted that the higher the MYC expression in a sample group, the more disorder would be observed. Using CCD, we indeed observed that C-MYC and MYC_SUM Quartile 4, which are the highest expression quartiles, were significantly disordered compared to quartile 2. However, it was also observed that in both cases MYC Quartile 1, the lowest expression quartile and lower than normal lung, was also significantly more disordered than Quartile 2 (**Figure 3D and E**).

Since the CCD results only reflected MYC expression and not activity, we next sought to determine if enrichment in the MYC targets pathway also correlated to circadian disorder, and in the same pattern. To make this comparison, GSVA was utilized to assign relative enrichment scores for the ‘Hallmarks MYC Targets V1’ pathway (42). Notably, this pathway consists of MYC downstream targets and represents MYC pathway activity rather than MYC expression itself. Using these relative enrichment scores for each LUAD sample, the samples were then sorted into quartiles and their CCD scores were compared. We found that the samples with the highest enrichment in the MYC pathway were the most disordered (**Figure 3F**). This agreed with the observation that those samples with the highest MYC expression also had disordered CCD scores. The fact that enrichment in the MYC pathway linearly correlated with increasing circadian disorder (ie, the lowest MYC pathway activation had the most ordered circadian rhythms) led us to hypothesize that the increased disorder in MYC expression quartile 1, where MYC was lower than normal lung, was due to a previously unknown and MYC-independent factor or pathway influencing the clock in cancer.

### EMT status is a MYC-independent factor that correlates with circadian status in LUAD tumors

We sought to understand what biological factors marked those tumors in the MYC expression quartile 1 group that had disrupted rhythms. One possibility was that the different MYC groups had different levels of tumor purity (percent of cancer cells as compared to percent of stromal and immune cells in the tumor). To determine if tumor purity could explain the difference in circadian disorder between MYC expression quartiles, we utilized cellularity scores for TCGA LUAD from a prior publication as a measure of tumor purity (36). We compared these scores based on MYC expression quartile and found that there was no significant difference in cellularity score across the MYC expression quartiles (**Supp.** Fig. 2A), suggesting that, on average, the tumors in each quartile had a similar balance of cancer and non-cancer cells. Interestingly, when cellularity score itself was used to group tumor samples, we found that high cellularity, which is high tumor purity, correlated with increased circadian disruption, and tumors with the lowest cellularity score (indicating less cancer cells compared to noncancer cells) had the most ordered rhythms (**Supp.** Fig. 2B). This agrees with a prior finding that solid tumors with higher expression of collagen genes (indicating more tumor-associated fibroblasts) had stronger circadian rhythms (43). These results indicated that while cellularity score itself correlated with circadian disorder, it does not explain the disorder in the lowest MYC expression quartile.

Another possibility was that the tumors in MYC quartile 1 were enriched for a different pathway associated with cancer. To identify the cause of the increased disorder in the lowest MYC expression quartile, we performed differential expression on MYC expression Quartiles 1 and 2 to determine which genes were differentially expressed between the two quartiles. We used CBioPortal to perform the differential expression analysis and found that there were thousands of genes that were differentially expressed between these groups (**Figure 4A**). We next performed Gene Set Enrichment Analysis (GSEA) to identify which pathways were enriched in each group. The higher MYC Quartile 2 (which had the lowest CCD score, corresponding with the most ordered clock function) had enrichment in pathways such as the Hallmarks MYC Targets Pathway V1, Hallmarks WNT Beta Catenin Signaling, TGF-Beta Signaling, and Epithelial to Mesenchymal Transition. The more disordered MYC Quartile 1 was enriched in KRAS Signaling Down, and CTNNB1 (β-catenin) Targets Down (**Figure 4B**).

**Figure 4:**
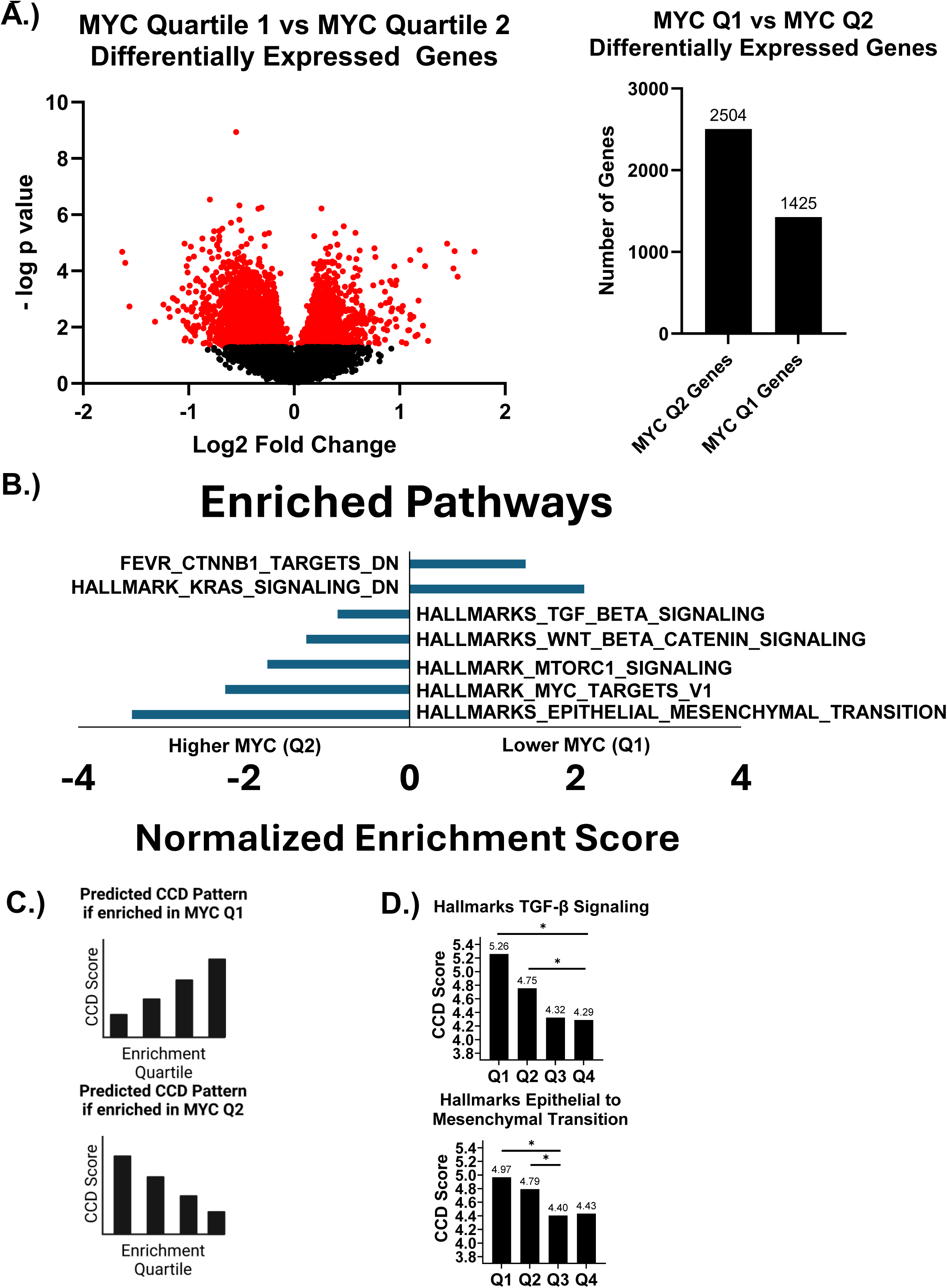
**Epithelial to mesenchymal transition and TGF-β Signaling negatively correlate to circadian disorder**. A.) Left Panel: A volcano plot shows the Log2 fold change of differentially expressed genes between the LUAD tumor samples in MYC_SUM quartile 1 and 2. All genes with a q-value below 0.05 are colored red. Right Panel: A bar chart representing the number of genes significantly differentially expressed between the two quartiles. B.) A bar chart of normalized enrichment scores for significantly differently enriched pathways identified through GSEA between MYC expression quartiles 1 and 2. C.) Predicted patterns of CCD scores if the tested pathway could be correlated with the disruption observed in MYC quartile 1 based on which MYC quartile the pathway is enriched in. D.) Bar charts of Hallmarks TGF-β Signaling and Hallmarks Epithelial to Mesenchymal Transition CCD scores based on enrichment quartiles of the respective pathway. Significance was determined by sample label permutation performed by the deltaCCD package. * = p<0.05.

We next sought to determine which of these differentially enriched pathways correlated to circadian disorder outside of the context of MYC. To test the connection between these pathways and circadian disorder in LUAD samples, GSVA was used to assign relative enrichment scores for each of the respective pathways for each tumor sample. These enrichment scores were used to sort the samples into new enrichment quartiles, based on relative enrichment of each pathway, and compared using CCD. We predicted that if a pathway correlated with the disruption that was observed in the MYC Quartile 1 expression quartile, its CCD scores would have a distinct pattern based on which expression quartile the pathway is enriched in (**illustrated in Figure 4C**). If the pathway was enriched in the MYC Quartile 1 and correlated with the observed disruption, then high enrichment in that pathway would be correlated with greater disruption, whereas pathways enriched in MYC Quartile 2 would have low enrichment quartiles correlated with greater disruption.

Two pathways emerged that matched the predicted pattern of enrichment and CCD score. Hallmarks Epithelial to Mesenchymal Transition (EMT) and TGF-β signaling pathways were enriched in MYC expression Quartile 2, and low enrichment in both pathways correlated to a significant increase in circadian disorder (**Figure 4D**). Since TGF-β can induce EMT in cancer cells, it logically followed that these two pathways showed a similar pattern. In contrast, while some of the other enriched pathways such as mTORC1 and KRAS showed correlation with circadian function, none matched the predicted pattern illustrated in **Figure 4C** that would explain disruption in MYC Quartile 1 (**Supp Figure 3**). This indicated that these two pathways could be correlated to the disorder observed in MYC expression Quartile 1.

### TGF-B treatment increases amplitude of oscillations in CMT64 cells *in vitro*

The observation that EMT and TGF-β signaling correlated with alterations in circadian rhythm were intriguing, as they suggested that more mesenchymal tumors may have stronger / more ordered rhythms. To directly test this potential correlation, we induced EMT *in vitro* with TGF-β in the CMT64 lung cancer cell line that we had transduced with a *BMAL1*::Luciferase reporter to track circadian rhythms with real-time luminescence monitoring (38). CMT64 cells were chosen for this study over the previously used cell lines, H1299 and A549, as they have robust rhythms. Previous studies using CMT64 cells with TGF-β showed that 5ng/mL TGF-β can induce morphological changes as well as induction of EMT (44). Upon treatment with 5ng/mL TGF-β for 72 hours, we observed that CMT64 cells became more elongated and spindle-like, exhibiting a mesenchymal-like phenotype compared to the control CMT64 cells (**Figure 5A**). We tested the expression of SNAIL in CMT64 control and TGF-β treated cells after 24 and 72 hours of exposure by qPCR and found that the expression of SNAIL was significantly increased in the treated cells after 72 hours (**Figure 5B**). Additionally, morphological changes induced by TGF-β treatment were further assessed by quantifying a roughness score, calculated from pixel intensity variation in brightfield images. This serves as a surrogate marker for structural changes occurring during epithelial to mesenchymal transition (**Figure 5C)**. In contrast, while E-Cadherin is often lost in cells undergoing EMT, we found that that E-Cadherin levels did not vary across the treatment conditions (**Figure 5D**) indicating that TGF-β potentially induced an intermediate ‘quasi-mesenchymal’ state in these cells(45). To test the role of TGFβ treatment in *BMAL1::*Luc oscillation, CMT64 cells were exposed to either vehicle control or 5ng/mL TGF-β for 72 hours before being placed in a Lumicycle live-cell luminometer. The TGF-β treated samples had a significant increase in the amplitude of their oscillation compared to control samples (**Figure 5E**), with no significant change in phase or dampening of their rhythms (**not shown**). These findings agree with our computational analysis correlating EMT and TGF-β signaling enrichment with more ordered rhythms, indicating that TGF-β and a mesenchymal state may directly modulate the circadian clock in lung adenocarcinoma.

**Figure 5:**
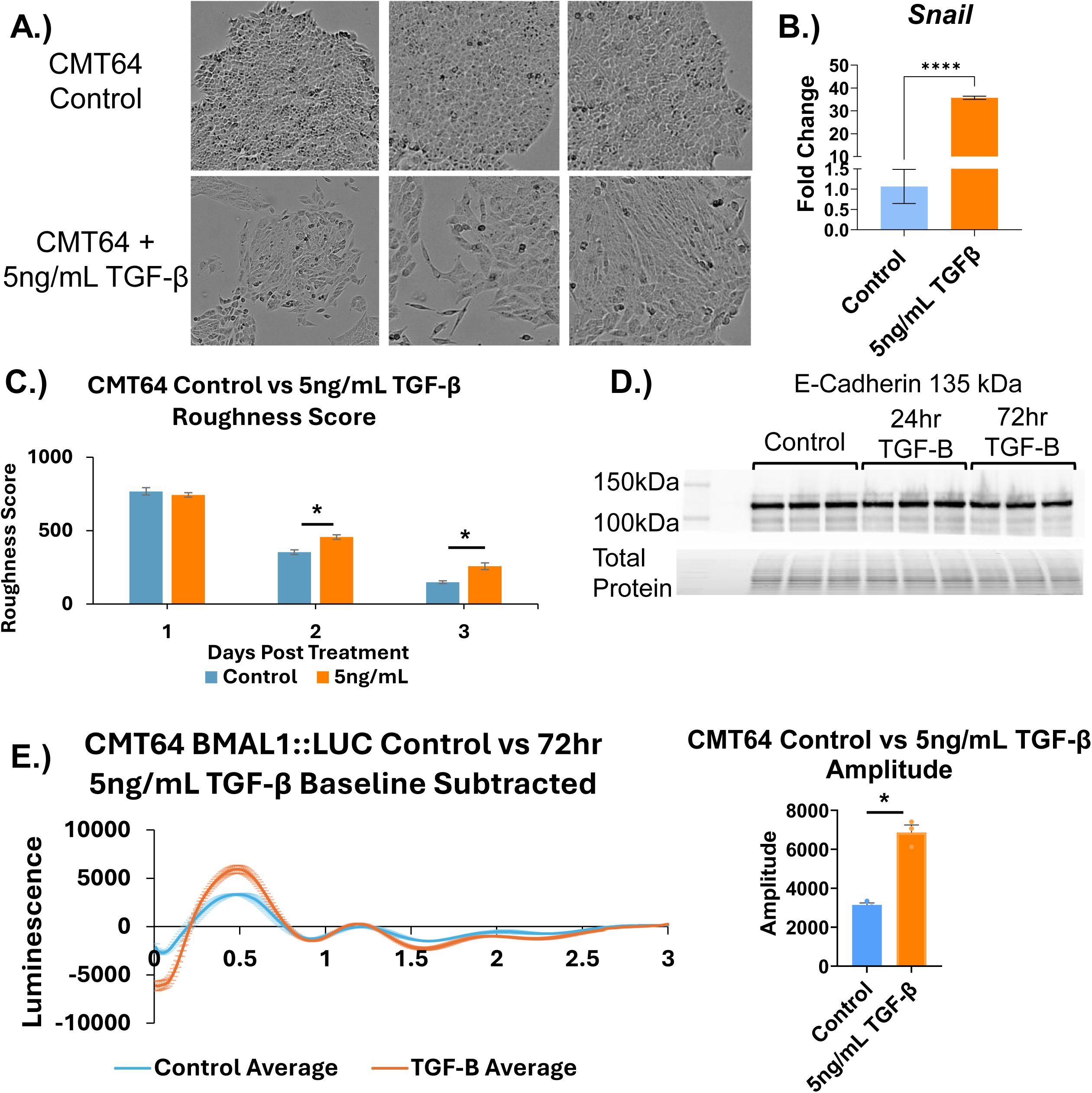
Treatment of CMT64 LUAD cells with TGF-β induces partial EMT and strengthens oscillations of BMAL1. A.) Top Row: Representative images of control CMT64. Bottom Row: Representative images of CMT64 cells that have been treated with 5ng/mL of TGF-β for 72 hours. B.) A bar chart of fold change in EMT marker SNAIL expression measured through qPCR in control CMT64 cells compared to cells exposed to 5ng/mL TGF-β for 72 hours with TBP used as a housekeeping gene. Significance was determined through Welch’s T-test. **** = p<0.0001. Error bars represent standard error of the mean. C.) A bar chart representing average roughness scores of control CMT64 cells and CMT64 cells treated with 5ng/mL TGF-β for 1, 2, and 3 days following treatment. Significance was determined by Welch’s T-test. * = p<0.05. Error bars represent standard error of the mean. D.) Western blot showing protein levels of E-Cadherin in CMT64 cells that were either control, treated with 5ng/mL TGF-β for 24 hours, or 5ng/mL TGF-β for 72 hours, compared with total protein as a loading control. E.) Lumicycle analysis of control CMT64 BMAL::LUC cells compared to those exposed to 5ng/mL TGF-β for 72 hours prior to being placed in the lumicycle. Oscillations are represented by plotting the luminescence over time with error bars representing the standard error of the mean. Calculated average amplitude is represented in a bar chart and significance was determined using a Welch’s T-test. * = p<0.05. Error bars represent standard error of the mean. For (B-E), N = 3 biological replicates and data are representative of 2-3 independent experiments.

## Discussion

To identify biomarkers of circadian disruption in lung cancer, we began by showing that NSCLC tumors have a more disordered clock compared to normal tissue samples. We then found that while the expression of some molecular clock genes correlate to survival, individual circadian gene expression is not a reliable marker for disorder in lung cancer. However, high and low MYC expressing tumors were significantly disordered compared to other tumors, and high enrichment in the MYC targets pathway correlated to circadian disruption. This did not explain the disruption observed in the low MYC group, which was lower than normal lung. To address this, we performed pathway analysis comparing the lowest MYC-expressing group, and the next highest group (which had the most ordered molecular clock scores from CCD). We then took the list of significantly differentially enriched pathways between the two MYC quartiles and ran Gene Set Variation Analysis to compare CCD score relative to enrichment of pathways associated with differential expression.

This analysis showed that two pathways (Hallmarks of Cancer Epithelial to Mesenchymal Transition and TGF-β signaling pathways) significantly correlated to disruption, and their quartile CCD scores matched the predicted pattern of a pathway that could explain the disruption in the low MYC tumors. To test this finding *in vitro*, mouse epithelial lung cancer cells were treated with TGF-β to induce EMT, and this treatment caused a significant increase in the amplitude of oscillations. These results indicate that MYC expression, MYC Target Pathway, EMT, and TGF-β Signaling pathway enrichment all correlate to circadian disorder in lung adenocarcinoma and could potentially be used as biomarkers for disruption in patients.

Our study suggests that gene pathways may be more reliable and consistent markers of circadian disruption in cancer than the expression of individual genes. For many individual genes that are part of the core circadian clock, such as *BMAL1*, there was no significant difference in circadian disorder based on their expression, or for some genes like *NR1D1* (REV-ERBα) the pattern of disruption correlated to their expression did not match with expectations based on their arm and function in the clock. Using pathway signatures appears to be more suited at identifying these factors that have significant impacts on the clock. When using a pathway signature for the circadian clock as a whole, as opposed to individual genes, we were able to determine that while there was high variance in the lower enrichment groups, the least enriched quartiles were more disordered than the most enriched samples. One potential way to improve this could be to refine the circadian gene list and consider the opposing positive and negative arms of the clock in building a pathway to use for GSVA.

Of the pathways that we identified as potential markers of circadian disruption, EMT and TGF-β pathways both individually correlate with circadian disruption outside of the context of MYC. We found that the high enrichment in these pathways was correlated to a stronger rhythm *in vitro* with our TGF-β treatment experiments showing a significant increase in the amplitude of oscillations. This pattern has previously been observed in cultures of human U20S cells, where disruption of TGF-β signaling caused the oscillations of the cells to decrease in amplitude and desynchronize from their neighbors(46). In that study, the Authors showed that TGF-β secretion coordinates and strengthens rhythms amongst neighboring cells in a paracrine fashion; however, EMT was not tested since the cells used were not epithelial in nature. Our method of treatment did not cause a full transition to a mesenchymal state, as the cells underwent some changes in morphology and SNAIL induction, but did not lose the epithelial cell marker E-Cadherin. This led us to the conclusion that these cells are better described as quasi-mesenchymal(45), and further experiments would be needed to determine if fully mesenchymal cells would have a greater change in their rhythms compared to untreated/epithelial cells.

These results indicate that low enrichment in EMT and TGF-β signaling pathway could indicate circadian disorder in human samples. Others have also observed relationships between EMT signaling and the molecular clock. A previous study, utilizing a method known as CYCLOPS which orders rhythms in untimed human data, showed that several genes that make up the EMT pathway have strong oscillations in breast cancer tumors with functioning clocks (17). This result combined with our own points towards a potential bidirectional relationship between the clock and EMT: when comparing tumors of high and low CYCLOPS magnitude, a measure of clock strength, EMT genes were identified as having the most change in their oscillations, while our results suggest that EMT enrichment in tumors can strengthen the clock in those samples. Together these findings suggest that the EMT pathway and the molecular clock could influence each other. Notably, the observation that more mesenchymal cells have stronger rhythms is counter to the simplified notion that clock disruption causes more aggressive cancer, and which may indicate that the clock is influencing the invasive potential of these tumors by promoting EMT at a later stage of cancer progression.

This could indicate that the role of the molecular clock shifts in cancer from being tumor-suppressive in early cancer development to pro-cancer in later stages. Our study shows that enrichment in the EMT and TGF-β pathways both correlate to clock strength through CCD analysis, and we were able to show that when epithelial mouse lung cancer cells were pushed to a quasi-mesenchymal state by TGF-β exposure they did have a significant increase in the amplitude of their rhythms confirming the effect.

We chose CCD for this analysis because it does not require time-series training data for new analyses and can produce data with a relatively small sample size. There have been other computational methods developed to interrogate single time point data for circadian analysis. The previously mentioned CYCLOPS method fits unordered data with no time labelling into a periodic cycle using global descriptors of gene expression and a machine learning algorithm, and can be used to identify cycling genes (12). Another computational method called LTM uses the observed variations in gene expression in human population data to identify genes that can influence the clock regardless of if they are rhythmic themselves (43). Finally, the TimeTeller method is a machine learning algorithm that instead of scoring the individual relationships between circadian genes like CCD, analyzes the clock as the multigenic system it is, allowing it to model the state of the clock based off the entire system and estimate overall clock function (11). TimeTeller requires substantial training data from normal samples, but this method could be used in the future to further explore the results of this study showing the impact of variations in enrichment of the MYC, EMT, and TGF-β pathways, and also points towards the importance of using the entire circadian system as a whole instead of individual genes to determine clock strength.

Overall, our results show that different factors, including MYC expression, MYC pathway activation, and EMT status, all correlate with the state of the molecular circadian clock in lung adenocarcinoma, and TGF-β directly influences the clock in lung adenocarcinoma cell lines. Our findings suggest that, in the absence of a direct measure of the strength of circadian rhythms in individual tumor samples, these pathways and potentially others could serve as indicators of circadian status in LUAD.

## Limitations of the study

One of the limitations of CCD as a process is its reliance on group level analysis and relationships between the data that is present in the study. While there are other computational methods to assess clock strength, such as the previously mentioned CYCLOPS, or assay the effect of genes on clock strength such as LTM, there is not currently a widely accepted method to determine and compare circadian strength at the single sample level, which would allow for a much greater understanding of the relationships between the clock and these varying biological factors that influence its rhythms. Also, these analyses only indicate correlations between these genes and pathways and circadian disruption. We will need further study and experiments to test the directionality of these correlations and to discover the mechanisms behind these relationships.

## Acknowledgments

The data and results shown here are in part based upon data generated by the TCGA Research Network: https://www.cancer.gov/tcga and the National Cancer Institute Clinical Proteomic Tumor Analysis Consortium (CPTAC).

## Author Contributions

**Jamison Burchett:** Conceptualization, Data Curation, Formal Analysis, Investigation, Methodology, Project Administration, Software, Validation, Visualization, Writing-original draft. **Aslihan Ambeskovic:** Methodology, Software. **McKayla Ford:** Methodology, Software. **Jacob Cody Naccarato:** Formal Analysis, Investigation, Validation, Visualization. **Juliana Cazarin:** Formal Analysis, Investigation, Resources, Validation, Visualization. **Fabio Hecht:** Methodology. **Molly Hulver:** Investigation. **Xueyang He:** Methodology, Software. **Joshua Munger:** Conceptualization. **Paula Vertino:** Supervision **Isaac Harris:** Supervision **Stephano Mello:** Conceptualization, Supervision **Brian J Altman:** Conceptualization, Data Curation, Funding Acquisition, Project Administration, Resources, Software, Supervision, Visualization, Writing-original draft.

**Supplemental Figure 1: Individual circadian genes have varied correlations to disorder.** A.) Expression of circadian genes is represented by violin plots of FPKM values with median expression represented by a dotted line, and significance determined by ordinary one-way ANOVA. * = p<0.05, ** = p<0.005, **** = p<0.0001. B.) Bar charts of the CCD score for each quartile of individual circadian genes with significance determined by the deltaCCD program. * = p<0.05. C.) A bar chart of CCD scores based on enrichment in a curated circadian gene set.

**Supplemental Figure 2: Tumor purity does not correlate with MYC expression but does correlate to circadian disorder**. A.) A violin plot of cellularity scores in each MYC expression quartile, with the median value represented with a dashed line. Significance was determined by ordinary one-way ANOVA. B.) A bar chart of CCD score for each quartile of tumor sample based on cellularity score with significance determined by the deltaCCD program. * = p<0.05.

**Supplemental Figure 3: Enrichment in biological pathways can be used as a marker for circadian disruption.** A.) CCD scores of enrichment quartiles of differentially enriched pathways are represented through bar graphs of the CCD score for each quartile with significance determined by the deltaCCD program. * = p<0.05.

