## Supplemental Figures and Legends for "MYC and Epithelial to Mesenchymal Transition (EMT) Independently Predict Circadian Rhythm Disruption in Lung Adenocarcinoma"

### Supplemental Figure 1

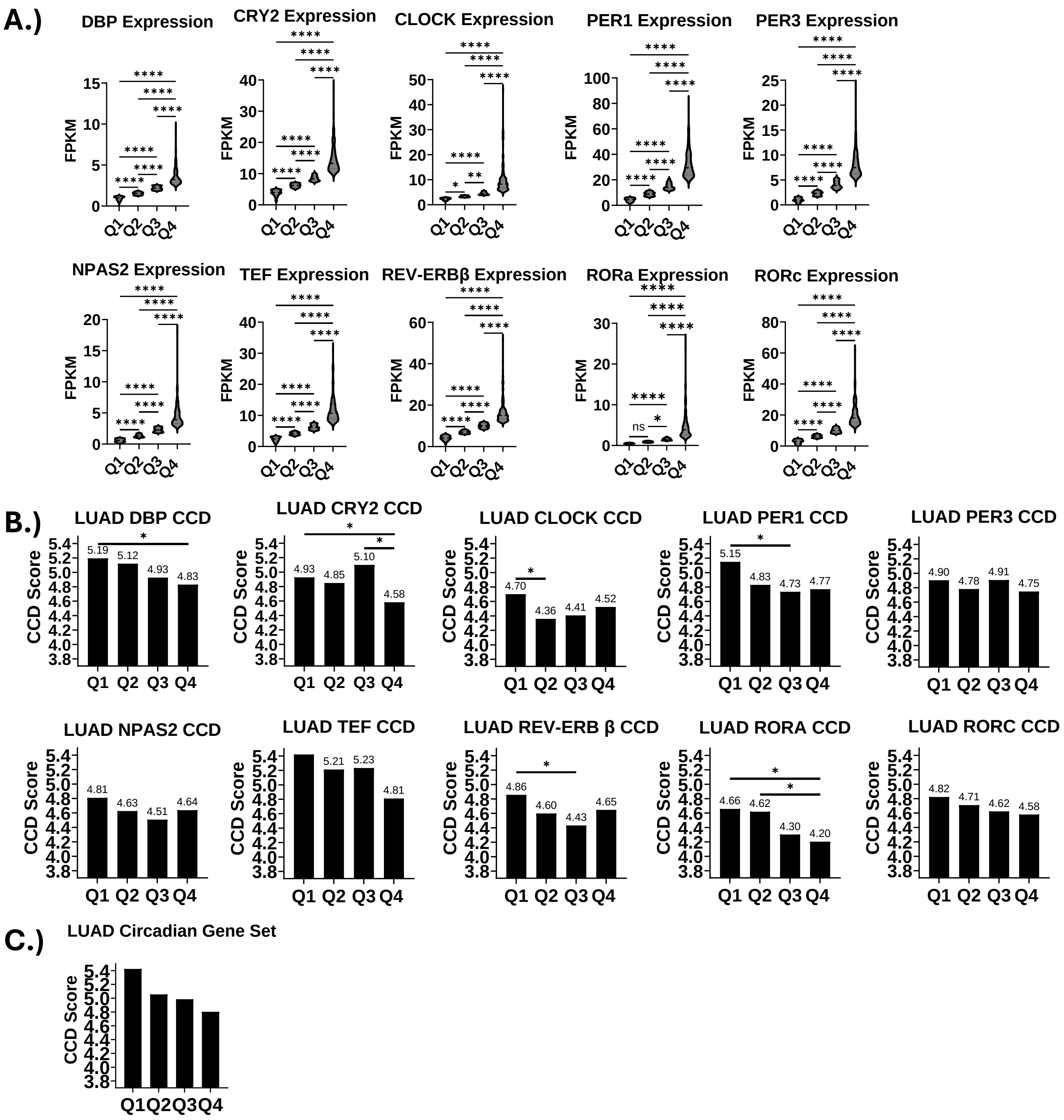


**Supplemental Figure 1: Individual circadian genes have varied correlations to disorder.** A.) Expression of circadian genes is represented by violin plots of FPKM values with median expression represented by a dotted line, and significance determined by ordinary one-way ANOVA. * = p<0.05, ** = p<0.005, **** = p<0.0001. B.) Bar charts of the CCD score for each quartile of individual circadian genes with significance determined by the deltaCCD program. * = p<0.05. C.) A bar chart of CCD scores based on enrichment in a curated circadian gene set.

Supplemental Figure 2

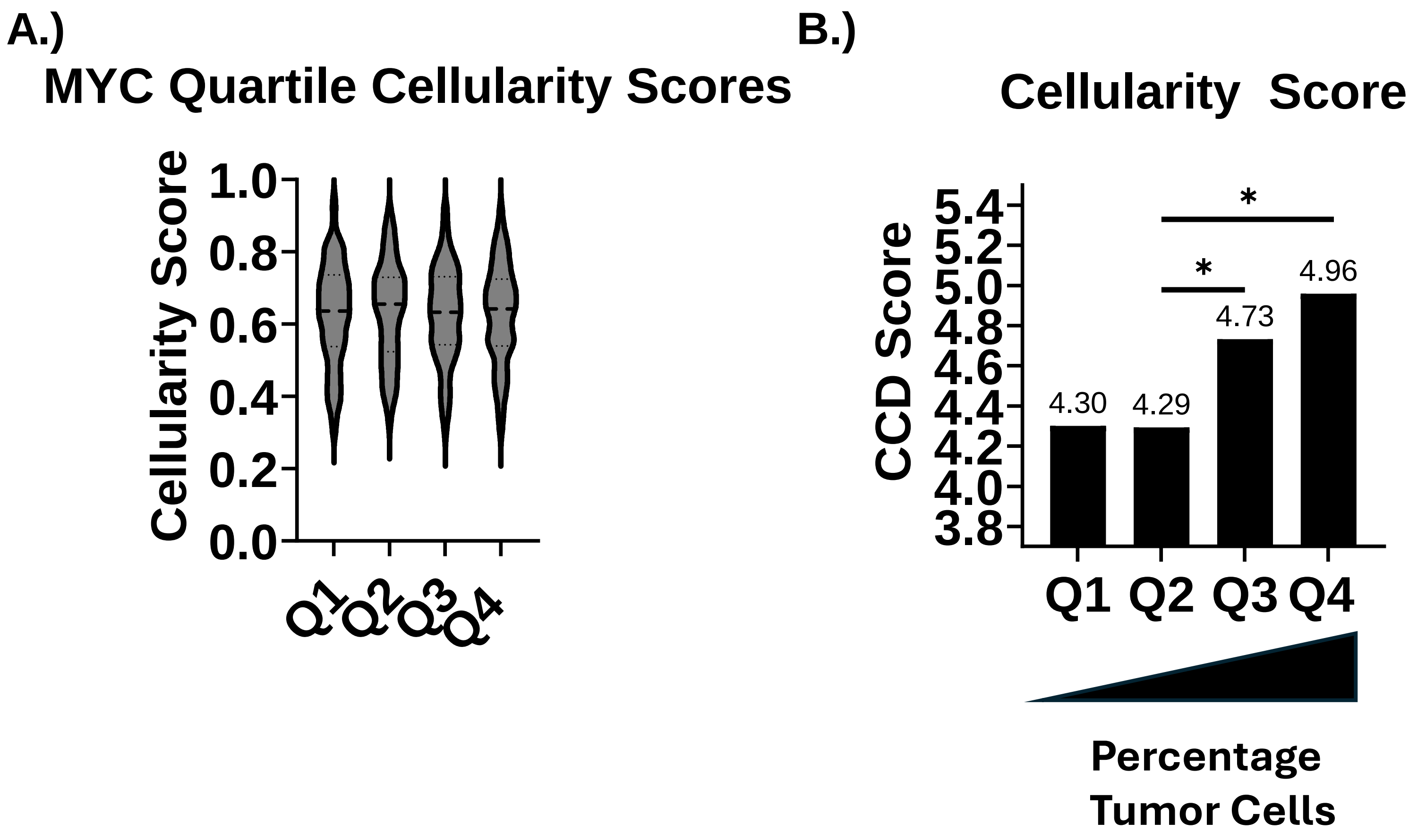


**Supplemental Figure 2: Tumor purity does not correlate with MYC expression but does correlate to circadian disorder**. A.) A violin plot of cellularity scores in each MYC expression quartile, with the median value represented with a dashed line. Significance was determined by ordinary one-way ANOVA. B.) A bar chart of CCD score for each quartile of tumor sample based on cellularity score with significance determined by the deltaCCD program. * = $p<0.05$.

### Supplemental Figure 3

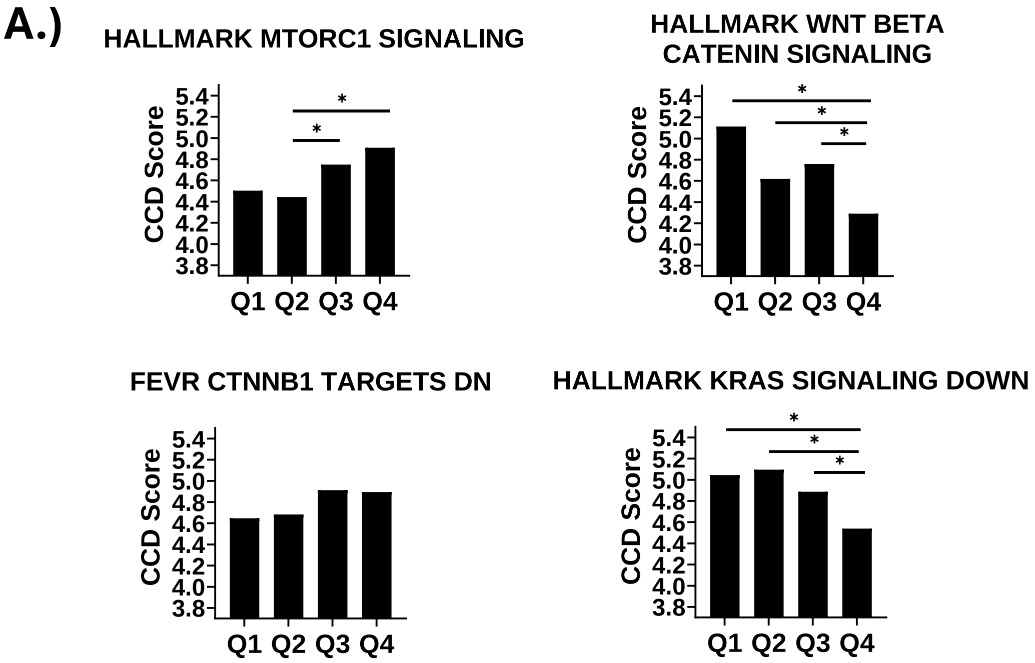


**Supplemental Figure 3: Enrichment in biological pathways can be used as a marker for circadian disruption.** A.) CCD scores of enrichment quartiles of differentially enriched pathways are represented through bar graphs of the CCD score for each quartile with significance determined by the deltaCCD program. * = p<0.05.
